# Haplotype assignment of longitudinal viral deep-sequencing data using co-variation of variant frequencies

**DOI:** 10.1101/444877

**Authors:** Juanita Pang, Cristina Venturini, Asif U. Tamuri, Sunando Roy, Judith Breuer, Richard A. Goldstein

**Affiliations:** Division of Infection and Immunity, University College London, London WC1E 6BT, United Kingdom; Research IT Services, University College London, London WC1E 6BT, United Kingdom; Infection, Immunity, Inflammation and Physiological Medicine, Institute of Child Health, University College London, London WC1E 6BT, United Kingdom; Great Ormond Street Hospital for Children, London WC1N 3JH, United Kingdom

**Author notes:** These authors contributed equally to this work (Juanita Pang, Cristina Venturini).

## Abstract

Longitudinal deep sequencing of viruses can provide detailed information about intra-host evolutionary dynamics including how viruses interact with and transmit between hosts. Many analyses require haplotype reconstruction, identifying which variants are co-located on the same genomic element. Most current methods to perform this reconstruction are based on a high density of variants and cannot perform this reconstruction for slowly evolving viruses. We present a new approach, HaROLD (HAplotype Reconstruction Of Longitudinal Deep sequencing data), which performs this reconstruction based on identifying co-varying variant frequencies using a probabilistic framework. We test this method with synthetic data sets of mixed cytomegalovirus and norovirus genomes, demonstrating high accuracy when longitudinal samples are available.

## INTRODUCTION

Next generation sequencing (NGS) of virus populations derived from medical and biological samples can deepen our understanding of virus biology, pathogen evolution, host-pathogen interactions, transmission dynamics and the development of drug resistance (Houldcroft, et al. 2017; Leung, et al. 2017; Moncla, et al. 2017). While smaller than bacterial and eukaryotic genomes, virus genomes are still much larger than the individual reads that are obtained through next generation sequencing. Detailed analyses often require determining which variants are found together in the same genome or genomic segment, a process known as haplotype reconstruction. This is commonly performed by identifying variants at sites that are close enough to be found on the same reads. If these variants are sufficiently dense, co-localised variants across the genome can be ‘stitched together’, resulting in the determination of whole genome haplotypes (Posada-Cespedes, et al. 2017). Unfortunately, viruses such as human cytomegalovirus (HCMV) can have long regions with few segregating sites, making it impossible to connect variants that span these regions.

There is increased focus on monitoring intra-host evolutionary dynamics using longitudinal sequencing, where samples are obtained from a single patient at multiple time points. Selection and drift result in changes in the relative frequencies of the haplotypes and thus in the frequencies of the variants that they contain. In such cases, we can use co-variation of variant frequencies to provide an additional source of information for haplotype reconstruction, even when these variants are far apart in the genome. In order to take advantage of this additional source of information, we created a new method for reconstructing whole-genome haplotypes from longitudinal sequence data (HAplotype Reconstruction Of Longitudinal Deep sequencing data, HaROLD), which was then used to analyse the high diversity of HCMV samples (Cudini, et al. 2019). We describe this method and demonstrate its utility and accuracy with synthetic data, both when multiple samples are available (as in the case for longitudinal sampling), as well as for single samples.

## RESULTS

### Evaluation based on synthetic data sets

In order to evaluate the ability of HaROLD to reconstruct haplotypes and estimate the relative haplotype frequencies, we created two sets of synthetic sequence data consisting of mixtures of whole genome sequences from GenBank, with multiple mixtures representing the results of longitudinal sampling. These data were broken into individual paired reads of length 250 and sequencing errors typical for Illumina sequencing were introduced. Set one consisted of mixtures of two to four norovirus sequences (approximately 7.5 kb in length), while the second set was assembled from two to three human cytomegalovirus (HCMV) sequences (approximately 230 kb). The sets of sequences ranged in similarity between 98.6% to 99.7% identity. The various synthetic sets are summarised in Tables 1 and 2.

**Table 1.**
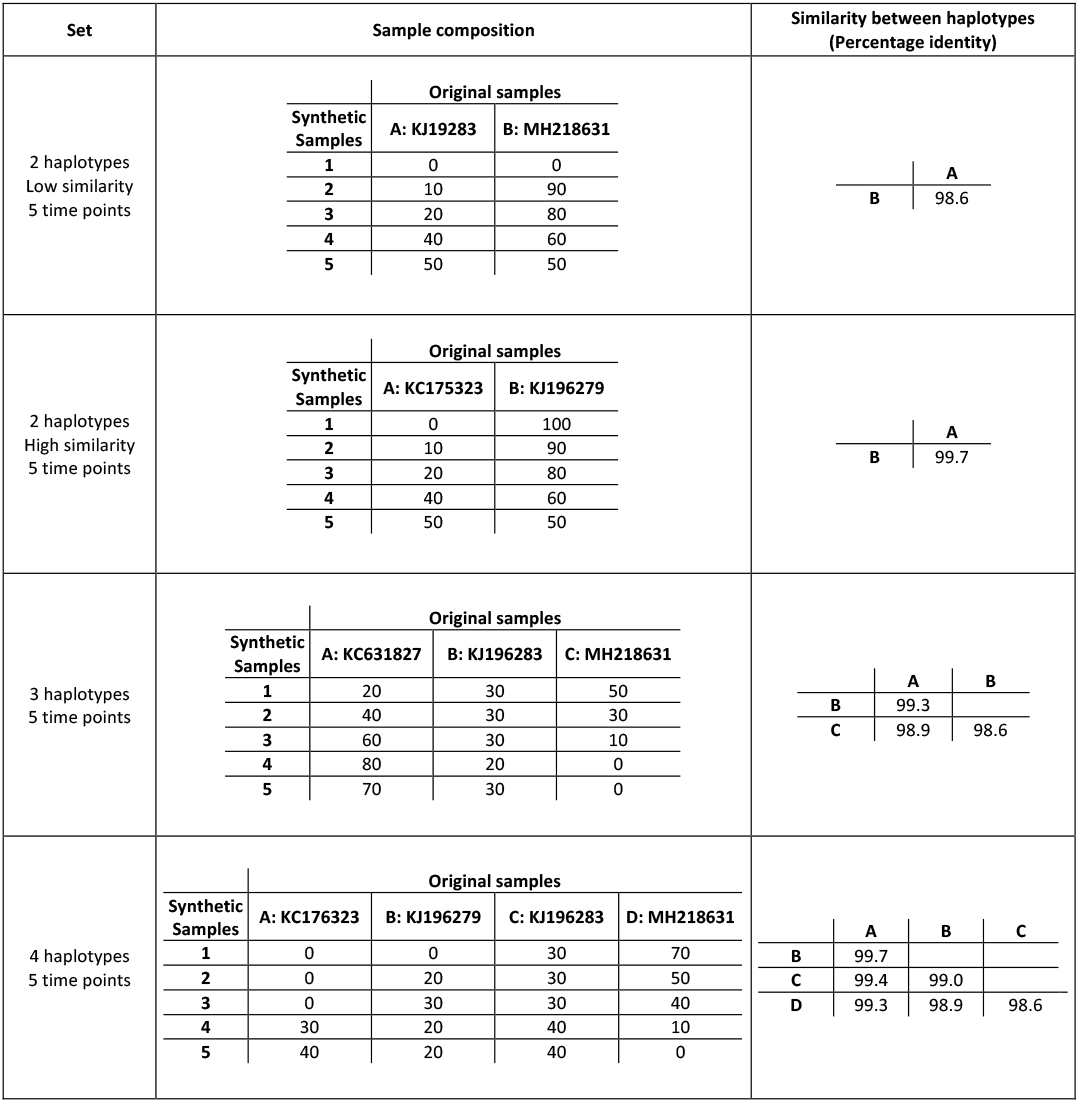
Summary of the longitudinal norovirus synthetic data sets used to test the accuracy of the haplotype reconstruction methods.

**Table 2.**
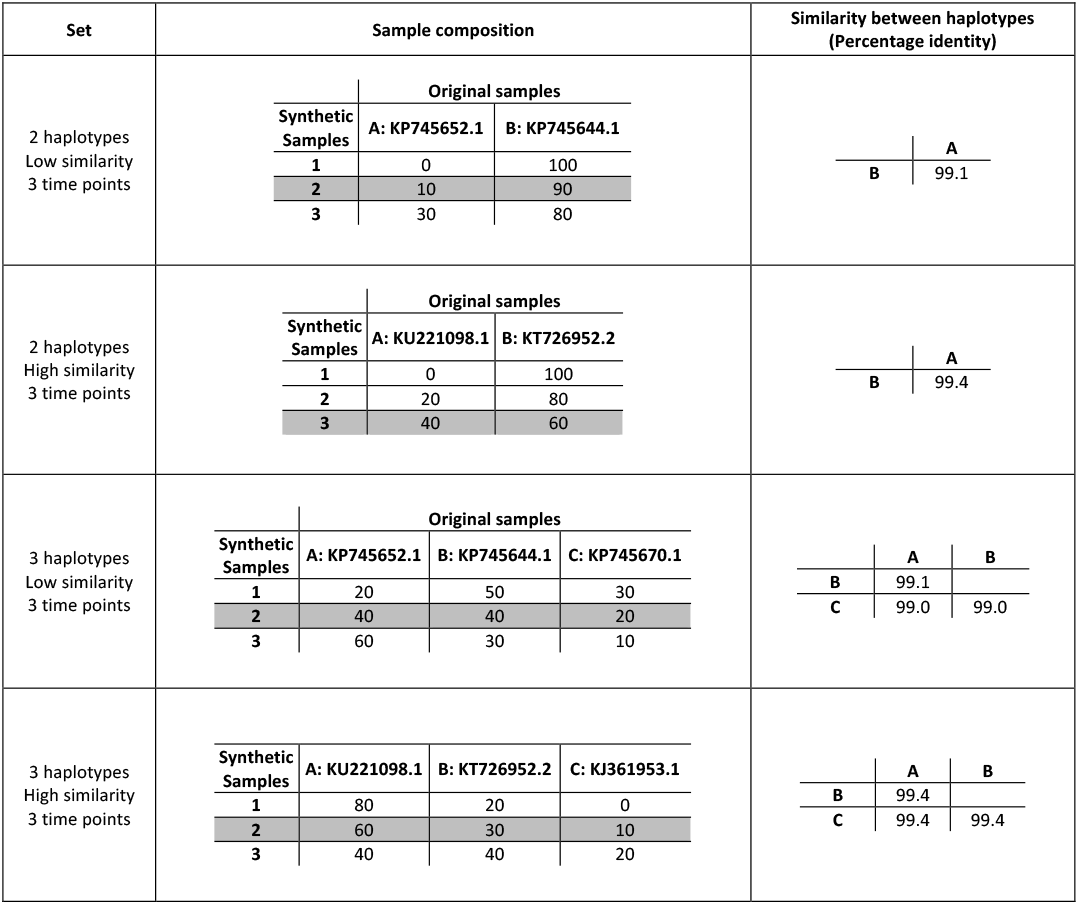
Summary of the longitudinal human cytomegalovirus synthetic data sets used to test the haplotype reconstruction methods. Samples highlighted in grey are used in the single time point test.

The performance of HaROLD for these data is represented in Figures 1 and 2. With the norovirus data, the reconstructed haplotypes were highly accurate, occasionally missing one or two bases at the end of the sequences (Figure 1A). The haplotype frequencies estimated by HaROLD were also highly accurate, with differences between the actual and estimated frequencies ranging from 0 to 0.002 (Figure 1B). Excellent results were also obtained with the synthetic data derived from HCMV; the reconstructed haplotypes were highly similar to the original sequences (similarity > 0.997) (Figure 2A) with differences between the actual and computed haplotype frequencies ranging from 0 to 0.06 (Figure 2B).

**Figure 1.**
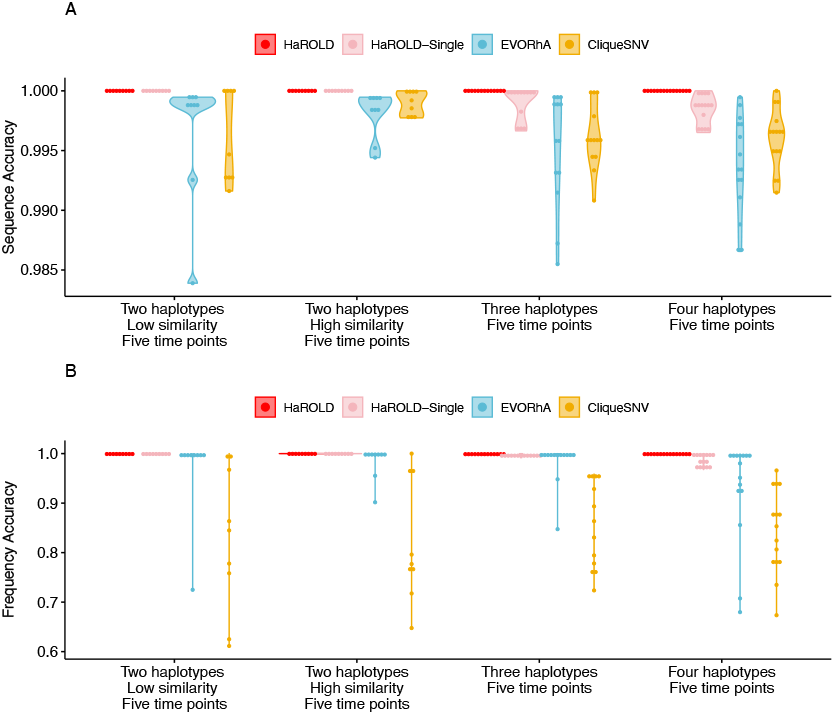
Violin plots showing the accuracy of haplotype reconstruction in the norovirus test set. (A) The accuracy of reconstructed sequence (pairwise distance between the actual sequence and reconstructed sequence). (B) The accuracy of computed frequencies (1 – |actual frequency – computed frequency|).

**Figure 2.**
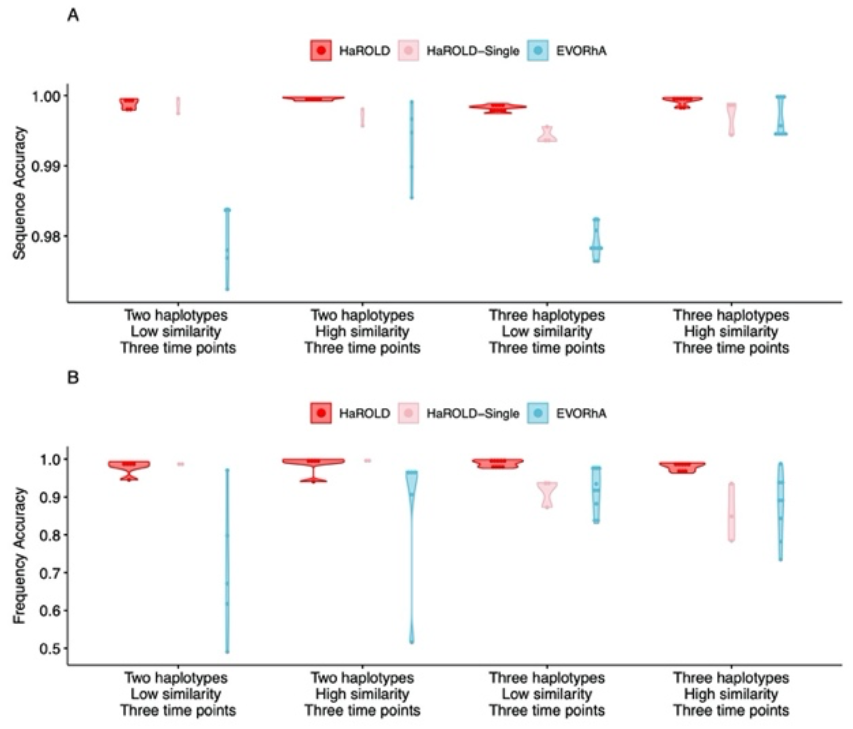
Violin plots showing the accuracy of haplotype reconstruction in the HCMV test set. (A) The accuracy of reconstructed sequence. (B) The accuracy of computed frequencies.

### Utility of longitudinal sampling

In contrast to most methods for haplotype reconstruction, HaROLD is formulated to take advantage of the availability of multiple longitudinal samples. In order to evaluate the importance of these longitudinal samples, we used HaROLD to reconstruct the haplotypes in our synthetic data sets without using this in additional information. This was accomplished by applying HaROLD to each sample independently, neglecting the presence of other related samples. As can be seen in Figures 1 and 2, the performance of HaROLD on single independent samples was generally worse, highlighting with the advantage of using longitudinal sampling information. Even so, the accuracies of the haplotype reconstructions were generally quite high, especially for the shorter norovirus sequences and when there were relatively few haplotypes

### Comparison with other methods

We compared the performance of HaROLD with two haplotype reconstruction techniques. The first method, CliqueSNV, constructs an allele graph based on linkage between variants (Knyazev 2020). The second method, EVORhA (Pulido-Tamayo, et al. 2015), is one of the few other haplotype reconstruction methods which also considers variant frequencies.

EVORhA generally estimated a larger number of haplotypes than present in the sample (ranging from 1 to 5 additional haplotypes), and consistently yielded haplotypes that most resembled the input reference sequence required for EVORha. The performance of EVORhA in estimating the relative haplotype frequencies was uneven.

On the norovirus datasets, CliqueSNV yielded more accurate haplotype sequences than EVORhA; frequency accuracy was, however, uneven. We were not able to successfully apply CliqueSNV to the HCMV datasets due to memory issues, despite the program running on an HPC node with greater than 70GB of memory).

In all these measures, HaROLD consistently outperformed these other techniques. Although the accuracy of the results declined when HaROLD ignored the presence of longitudinal samples and considered each sample independently, even in such cases the results obtained with HaROLD were generally equal or superior to these other two methods.

As an example of the consequences of the different reconstruction accuracies on downstream analyses, we estimated the diversity of the various samples based on the reconstructed haplotypes, as shown in Figure 3. The haplotypes generated by EVORhA and CliqueSNV generally produced exaggerated estimates of the within-sample diversity, with the exception of the two haplotypes test sets in norovirus where the haplotypes produced by EVORhA resulted in an underestimate. The haplotypes generated by HaROLD resulted in significantly better estimates of the diversity, especially when the reconstruction took advantage of the presence of longitudinal samples.

**Figure 3.**
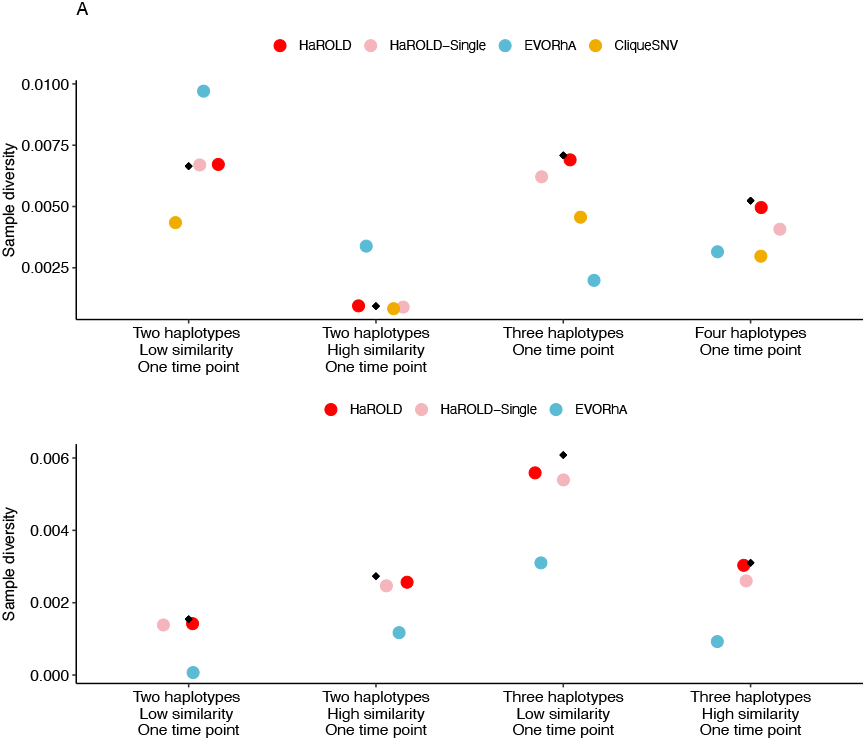
Accuracy of sample diversity estimations based on reconstructed haplotypes for norovirus test set (A) and HCMV test set (B). True sequence diversity shown with black diamonds.

## DISCUSSION

The majority of methods for reconstructing haplotypes rely on reads that contain multiple polymorphic sites, and thus require a sufficiently density of polymorphic sites so that the distances between such sites are closer together than the read length. Unfortunately, this is not always the case, especially for viruses such as HCMV where much of the observed sequence diversity is confined to short intervals between protein coding regions. Even when there is copious variation, there may be closely related haplotypes where the haplotype-defining variants are separated by distances greater than the read length, making it difficult to assign these variants correctly to the otherwise similar haplotypes. HaROLD was motivated by the increasing availability of multiple samples that are likely to share haplotypes, such as longitudinal studies of within host evolution or samples from an outbreak cluster. Under such conditions, variant frequencies can provide an important additional source of information for making accurate haplotype reconstructions. Notably, HaROLD generates haplotypes as accurate or more accurate than other tested methods even when multiple samples were not available. This greater accuracy was achieved with significantly less computing power and memory than the other methods we used for comparison, allowing rapid analysis of sequence data, even for bigger DNA viruses such as HCMV.

It is difficult to determine how many haplotypes there are in the sample, even with perfect information. One could consider every unique sequence in the sample as a different haplotype, but in this case the number of haplotypes would often be so large as to make any further analysis impractical. Alternatively, one could consider haplotypes as representing clusters of closely related sequences that do not need to be all exactly identical. In this case, there is some flexibility in how one defines the term ‘closely related’. HaROLD is generally conservative about the number of haplotypes. In particular, the refinement method does not add an additional haplotype unless the improvement in the log likelihood is sufficient to justify the resulting increase in the number of parameters. The resulting haplotypes then include some amount of variation, which is provided as output to the user. In particular, the output reports the probability that a sequence belonging to a haplotype would have any of the four bases found in each site. When these probabilities are sufficiently definitive, a base is assigned in the multiple sequence alignment. An ambiguous base is presented when a definitive assignment cannot be made.

## MATERIAL AND METHODS

We briefly describe the methods here. Further details are included in the Supplementary materials.

### Initial estimation

Consider a set of related samples that have been analysed using NGS. These may, for instance, represent a series of virus samples that have been extracted from a single patient at various time points. We initially assume that these samples contain a common set of haplotypes but in differing proportions, an assumption that will be relaxed at a later stage. Note that the number of samples can be as small as one, and each sample does not necessarily contain every haplotype.

We start with an assumed total number of haplotypes for the set of samples. Following quality control and assembly of the reads, for each sample we count the number of each type of base observed at each position in the resulting alignment. The observed number of each base depends on a) the frequencies of the haplotypes in that sample, b) the base found at that position in each of the haplotypes, and c) the probability of making an erroneous measurement at that site. As the error rate may be different at different sites and on different strands, we consider that this rate is drawn from a Dirichlet distribution. We first find the maximum likelihood estimate of the haplotype frequencies in each sample and the parameters defining the error rate distribution. We account for our initial ignorance of the haplotype sequences by summing this likelihood over all possible ways the different bases observed at that position can occur in the different haplotypes; if three different bases are observed at that position and our current model involves five haplotypes, we sum over all 3^5^ = 243 possible assignments of bases to haplotypes. We also integrate over the distribution of error rates. In this manner, the sequences of the haplotypes and the site- and strand-specific error rates initially represent nuisance parameters.

Following estimation of the haplotype frequencies and error rate distribution parameters, we determine how much each assignment of bases to haplotypes contributes to the overall likelihood. This allows us to calculate the posterior probability of each assignment of bases to haplotypes. By summing over these posterior probabilities, we can compute the marginal posterior probability that a base is found at that site in each of the haplotypes. If these probabilities are sufficiently definitive, an assignment is made. The *a posteriori* marginal probability of each base is included in the output.

We perform this procedure for a range of different numbers of haplotypes. As increasing the number of haplotypes increases the number of ways of assigning bases to each of the haplotypes, decreasing the prior probability of any given assignment, the log likelihood typically decreases when the number of haplotypes increases beyond that necessary to represent the data. We select the number of haplotypes that maximise the log likelihood.

The run time is strongly dependent on the number of haplotypes, average read depth and size of the virus; run times for the synthetic data described in Tables 1 and 2 (7.5kb, 1 to 4 haplotypes and 235kb, 1 to 3 haplotypes) ranged from 6s to 30m on a single HPC node (8G). This can be longer in some cases, generally dominated by the estimation of the error rate parameters. The calculations can, correspondingly, be greatly sped up if these parameters are estimated and fixed. HaROLD supports multiple threads.

### Further refinement

The method described above takes advantage of the presence of the same haplotype in multiple samples at various frequencies. It assumes that these haplotypes are identical in the various samples, neglecting processes such as mutations. It also ignores the information that forms the basis of most haplotype reconstruction methods, the presence of multiple variants on the same read. The next step is to relax these assumptions and use variant co-localisation to refine the haplotypes.

For this next stage, each sample is analysed individually. We start with the estimated frequencies of each haplotype in this sample, and the *a posteriori* probability of each base at each site in each haplotype, as output from the previous program. The haplotypes are then optimised by assigning the reads, probabilistically, to the various haplotypes. The number of reads assigned to each haplotype is used to adjust the frequencies of each haplotype. The reads are then re-assigned until the haplotype frequencies have converged. The resulting assigned reads are then used to update the probability of the bases found in each site in all of the reads assigned to each haplotype. This process of is performed until convergence.

If requested by the users, a number of modifications of the haplotypes are considered. These include a) recombination of two haplotypes, where corresponding regions of the haplotype sequences are swapped, b) gene conversion, where a region of one haplotype sequence is overwritten by the corresponding region of a different haplotype sequence, c) merging of two haplotypes into a single new haplotype, reducing the total number of haplotypes by one, and d) dividing a single haplotype into two new haplotypes, increasing the total number of haplotypes by one. After each proposed modification, the haplotype sequences and frequencies are re-optimised until convergence. The modification is then rejected or accepted based on the comparison of the Akaike Information Criterion (AIC) (Akaike 1974) of this new set compared with the AIC of the set of haplotypes prior to the attempted modification.

## Supporting information

Details of methodology

## DATA AND SOFTWARE AVAILABILITY

The software HaROLD and other materials used is available in the GitHub repository https://github.com/ucl-pathgenomics/HAROLD

## SUPPLEMENTARY DATA

Supplementary Data are available at BioRχiv.

## ACKNOWLEDGEMENTS

Wellcome Trust Collaborative Awards [203268, 204870], Rosetrees Trust PhD Studentship [M876], Medical Research Council [MC_U117573805].

