## Supplementary material for "Haplotype assignment of longitudinal viral deep-sequencing data using co-variation of variant frequencies": Details of methodology

### SUPPLEMENTARY MATERIALS

### PREPARATION OF SYNTHETIC TEST DATA SETS

In order to evaluate the ability of HaROLD to reconstruct haplotypes and the relative haplotype frequencies, we created synthetic data sets consisting of mixtures of whole genome norovirus sequences from GenBank (Benson, et al. 2013), with multiple mixtures representing longitudinal sampling. The various synthetic sets are summarised in Table 1, involving various combinations of KC175323, KC631827, KJ196279, KJ196283 and MH218631.

SimSeq (Benidt and Nettleton 2015) was used to create 1,000,000 paired end reads of length 250 for each GenBank sequence listed in Table 1 and Table 2. The output SAM files from SimSeq were then converted into Fastq files using Picard version 2.21.1 'SamToFastq' (Institute 2019). In order to construct the data sets, Seqtk (Shen, et al. 2016) was used to subsample and mix the reads from each ensemble according to the relative fractions listed in Tables 1 and 2. Reads were then trimmed for adapters using Trim galore version 0.6.0 (Bioinformatics 2019). Duplicate reads were removed using Picard version 2.21.1 'MarkDuplicates' (Institute 2019). Reads were mapped to a norovirus GII.Pe-GII.4 Sydney 2012 reference strain JX459907 using BWA version 0.7.17 (Li and Durbin 2009). The Makereadcount.jar (https://github.com/ucl-pathgenomics/HaROLD/tree/master/jar) was used to obtain the strand specific nucleotide counts from BAM files. These strand count files were used as the input for HaROLD.

We also created synthetic data sets consisting of mixtures of whole genome human cytomegalovirus (HCMV) sequences from GenBank, with the same method and in the same format as the norovirus data sets. The various data sets are summarised in Table 2, involving various combinations of KP745652.1, KP745644.1, KU221098.1, KP745670.1, KJ361953.1 and KT726952.2. We simulated 100,000 paired end reads with the same approach used for norovirus and mapped to reference strain Merlin (NC_006273.2), resulting in 80-90x average coverage and >99% of the genome covered.

### DETAILS OF THE METHOD

The haplotype analysis looks at matching a statistical model to longitudinal data of the form $\{n_{b,s,l,t}\}$ representing the number of reads with base $b$ on strand $s$ at position $l$ derived from the sample acquired at timepoint $t$. We consider that the reads come from a set of haplotypes where $\{x_{j,1}, x_{j,2}, x_{j,3} \dots x_{j,M}\}$ is the sequence of length $M$ of haplotype $j$. At time point $t$, we represent the frequencies of the $k$ different haplotypes by $\{\Pi_{1,t}, \Pi_{2,t}, \Pi_{3,t} \dots \Pi_{k,t}\}$ which obey $\sum_j \Pi_{j,t} = 1$. Multiple haplotypes might share the same base at a given location; the frequencies of base $b$ at site $l$ at timepoint $t$ is equal to the sum of the frequencies of all haplotypes that have that base at that site and is equal for the two strands: $\pi_{b,s,l,t} = \sum_{j \,\forall\, x_{j,l}=b} \Pi_{j,t}$.

#### Computing likelihoods

We consider that, for a site with an error rate of $\epsilon$ there is a probability $1 - 3\epsilon$ of observing the true base and a probability $\epsilon$ of observing one of the three other bases. If we know $\pi_{b,s,l,t}$, then the probability $P_{s,l,t}(b_i)$ of a read at that time, strand and position being observed as a base $b_i$ is given by

$$P_{s,l,t}(b_i) = \pi_{b_i,s,l,t}(1 - 3\,\epsilon) + \epsilon \sum_{b_j \neq b_i} \pi_{b_j,s,l,t} = \pi_{b_i,s,l,t} + \left(1 - 4\,\pi_{b_i,s,l,t}\right)\epsilon \qquad (1)$$

where we have used the fact that $\sum_b \pi_{b,s,l,t} = 1$. The probability of observing $\{n_{b,s,l,t}\}_{s,l,t}$ , the reads for a specified strand, site, and time, is then given by the multinomial distribution

$$P\left(\{n_{b,s,l,t}\}_{s,l,t} | \{\pi_{b,l,t}\}, \epsilon\right) = \frac{n_{s,l,t}!}{\prod_{b_i} n_{b_i,s,l,t}!} \prod_{b_i} \left(\pi_{b_i,s,l,t} + \left(1 - 4\,\pi_{b_i,s,l,t}\right)\epsilon\right)^{n_{b_i,s,l,t}} \tag{2}$$

We do not know the values of $\epsilon$, and cannot assume that these probabilities are the same for all strands, bases, and time points or bases. Rather, we describe a distribution of probabilities of the bases where Equation (2) is satisfied on average. We do this by constructing a Dirichlet distribution $\mathrm{Dir}_{\{\alpha_{b,s,l,t}\}}\left(\{P_{s,l,t}(b_i)\}\right)$ where $\alpha_{b_i,s,l,t} = \alpha_0\,\pi_{b_i,l,t} + \alpha_\epsilon$. For such a Dirichlet distribution, the average probability of observing a given base $\langle P_{s,l,t}(b_i)\rangle$ is given by

$$\langle P_{s,l,t}(b_i)\rangle = \frac{\alpha_0\,\pi_{b_i,l,t} + \alpha_\epsilon}{\alpha_0 + 4\alpha_\epsilon} = \pi_{b_i,l,t} + \left(1 - 4\,\pi_{b_i,l,t}\right)\left(\frac{\alpha_\epsilon}{\alpha_0 + 4\alpha_\epsilon}\right) \tag{3}$$

which mirrors Equation **Error! Reference source not found.** when $\epsilon = \frac{\alpha_\epsilon}{\alpha_0 + 4\alpha_\epsilon}$.

The advantage of this approach is that we are not specifying an error rate but allowing the error rate to vary by an amount determined by the parameters in the Dirichlet distribution, $\alpha_0$ and $\alpha_\epsilon$. We are assuming that the distribution is symmetric with respect to the various bases, but the rates for specific errors need not be the same. We now have a distribution of $\{P_{s,l,t}(b_i)\}$ rather than specific values, so in order to calculate the likelihood of the observed data on that strand at that position and time, we need to integrate over this distribution

$$\Lambda\left(\{n_{b,s,l,t}\}_{s,l,t} | \{\vec{x_j}\}_l, \{\Pi_{j,t}\}, \alpha_0, \alpha_\epsilon\right) = \frac{n_{s,l,t}!}{\prod_{b_i} n_{b_i,s,l,t}!} \int \prod_{b_i} \left(P_{s,l,t}(b_i)\right)^{n_{b_i,s,l,t}} \mathrm{Dir}_{\{\alpha_{b,s,l,t}\}}\left(\{P_{s,l,t}(b_i)\}\right)\, d\left(\{P_{s,l,t}(b_i)\}\right) \tag{4}$$

where the integral is over the entire range of the Dirichlet distribution, which is all values of $\{P_{s,l,t}(b_i)\}$ such that $\sum_i P_{s,l,t}(b_i) = 1$, and we have explicitly considered that the calculations of $\pi_{b_i,l,t}$, and thus $P_{s,l,t}(b_i)$, rely on $\{\vec{x_j}\}_l$, the haplotype sequences $\{\vec{x_j}\}_l$at position $l$ , as well as the haplotype frequencies and the parameters $\alpha_0$ and $\alpha_\epsilon$. Taking advantage of the properties of the Dirichlet distribution, this integral is quite easy, resulting in

$$\Lambda\left(\{n_{b,s,l,t}\}_{s,l,t} | \{\vec{x_j}\}_l, \{\Pi_{j,t}\}, \alpha_0, \alpha_\epsilon\right) = \frac{n_{s,l,t}!}{\prod_{b_i} n_{b_i,s,l,t}!} \frac{\mathrm{B}\left(\{\alpha_0\,\pi_{b_i,l,t} + \alpha_\epsilon + n_{b_i,s,l,t}\}\right)}{\mathrm{B}\left(\{\alpha_0\,\pi_{b_i,l,t} + \alpha_\epsilon\}\right)} \tag{5}$$

where $\mathrm{B}(\{\theta_i\})$ is the multivariate beta function defined as $\mathrm{B}(\{\theta_i\}) \equiv \frac{\Gamma(\theta_1)\Gamma(\theta_2)\ldots\Gamma(\theta_n)}{\Gamma(\theta_1+\theta_2+\cdots+\theta_n)}$.

**Distributing bases amongst haplotypes**

The calculation of $\pi_{b,s,l,t} = \sum_{j\,\forall\,x_{j,l}=b} \Pi_{j,t}$ relies on knowing the haplotype sequences $\{x_{j,1}, x_{j,2}, x_{j,3} \ldots x_{j,M}\}$. As this information is unavailable *a priori*, we calculate the likelihood for all $k^M$

possible ways of assigning $k$ bases to $M$ haplotypes, where $k$ is the number of unique bases observed at that position. The haplotype sequence is assumed to be the same for both strands and all time points; therefore, the sum over possible haplotype sequences is outside the sum over strands and time points, but we can consider each site separately.

$$\Lambda\left(\{n_{b,s,l,t}\}\right) = \prod_{l} \sum_{\{\vec{x_j}\}_l} \frac{1}{4^M} \prod_{s,t} \Lambda\left(\{n_{b,s,l,t}\}_{s,l,t} | \{\vec{x_j}\}_l, \{\Pi_{j,t}\}, \alpha_0, \alpha_\epsilon\right) \quad (6)$$

where the sum is over all possible distributions of bases amongst haplotypes at position $l$.

We first maximise this expression by adjusting the values of $\{\Pi_{j,t}\}$, $\alpha_0$, and $\alpha_\epsilon$, noting that $\{\Pi_{j,t}\}$ is the same for all locations at each time point. By considering which assignments of bases to haplotypes contribute the most to the likelihood, we are able to calculate posterior probabilities of the arrangement of bases at each position in the sequence, allowing us to calculate the posterior probability of each base at each position on each haplotype.

**Refining the haplotypes**

The method described above takes advantage of the presence of the same haplotype in multiple samples at various frequencies. It assumes that these haplotypes are identical in the various samples, neglecting processes such as mutations and recombination events. It also ignores the information that forms the basis of most haplotype reconstruction methods, the presence of multiple variants on the same read. The next step is to relax these assumptions and use the co-variation to refine the haplotypes.

For this next stage, each sample is analysed individually. A flowchart of the refinement process is shown in Supplementary Figure 1. We start with the estimated frequencies of each haplotype $j$ in this sample, $\{\Pi_j\}$, and a posteriori probability of each base $b_i$ at each site l in each haplotype, $\pi_{j,l}(b_i)$, as output from the previous program. The haplotypes are then optimised by assigning the reads, probabilistically, to the various haplotypes. The number of reads assigned to each haplotype is used to adjust the frequencies of each haplotype. The reads are then re-assigned until the haplotype frequencies have converged. The resulting assigned reads are then used to update $\pi_{j,l}(b_i)$ based on the bases found in each site in all of the reads assigned to each haplotype. This process of updating $\{\Pi_j\}$ and $\pi_{j,l}(b_i)$ is performed until convergence.

The next step, if requested by the user, is to consider an adjustable number of possible recombination of the haplotypes. These recombination events involve a) picking two haplotypes at random, b) picking a region of the alignment, of length chosen from a Normal distribution with standard deviation of 10 sites, and then c) either swapping the values of $\pi_{j,l}(b_i)$ in this region between the two haplotypes (50% probability) or over-writing the values in one haplotype with the values from the other (25% probability for each direction.) Following such a step, the haplotype frequencies and base probabilities are then re-optimised as described above, and the recombination event is either accepted or rejected based on whether the penalised log likelihood, that is, the log likelihood minus the number of adjustable parameters defining the haplotypes, is increased or decreased.

If requested by the user, the program then implements an iterative process of refinement. At the start of each iteration, if requested, pairs of haplotypes are chosen and merged, with the frequencies of the resulting haplotype equal to the sum of that of the parents, and the base frequencies equal to the average of the two parents. This results in a reduction in the number of haplotypes by one. The

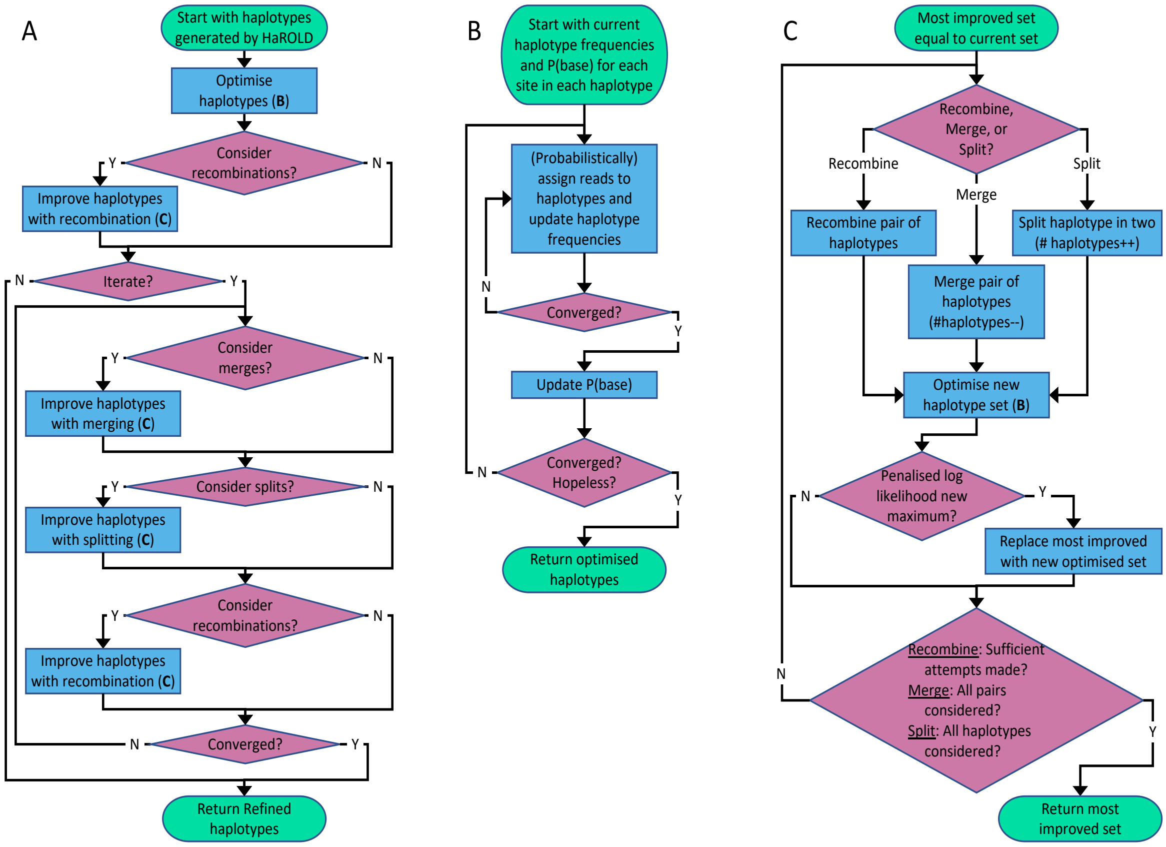


**Supplementary Figure 1. Flowchart of refinement process.** (A) Overall process. (B) Subprocess for haplotype optimisation. (C) Subprocess for considering recombination, merging and splitting; merging decreases the number of haplotypes by one, whilst splitting increases this number by one. Support for these three operations is evaluated by considering the Akaike Information Criterion (AIC). Following optimisation; the number of adjustable parameters is estimated by the number of haplotypes times the number of different bases found at each site minus one, summed over all sites.

haplotypes are then re-optimised. This process is repeated for every pair of haplotypes. The merge that most increases the penalised log likelihood is recorded.

Next, if requested, a haplotype is chosen and split into two haplotypes, increasing the total number of haplotypes by one. The resulting set of haplotypes is then re-optimised. This is repeated for every original haplotype. The split that results in the largest increase in penalised log likelihood is recorded.

Finally, if requested, the recombination process described above is performed. Again, the recombination event that results in the largest increase in penalised log likelihood is recorded. Following these attempted modifications of the haplotypes, the modification – merge, split, or recombination – that most increases the penalised log likelihood is compared with the penalised log likelihood at the beginning of the iteration. If this results in a net increase in the penalised log likelihood, this modification is accepted, and becomes the starting position for the next iteration. This iterative process is then repeated until convergence.
